# Impaired potency of neutralizing antibodies against cell-cell fusion mediated by SARS-CoV-2

**DOI:** 10.1101/2023.03.09.531948

**Authors:** Qian Wang, Andre Yanchen Yeh, Yicheng Guo, Hiroshi Mohri, Jian Yu, David D. Ho, Lihong Liu

**Affiliations:** Aaron Diamond AIDS Research Center, Columbia University Vagelos College of Physicians and Surgeons, New York, USA; School of Medicine, National Taiwan University, Taipei, Taiwan; Department of Microbiology and Immunology, Columbia University Vagelos College of Physicians and Surgeons, New York, NY, USA; Division of Infectious Diseases, Department of Medicine, Columbia University Vagelos College of Physicians and Surgeons, New York, NY, USA

**Keywords:** SARS-CoV-2, Omicron subvariants, spike protein, cell-cell fusion, monoclonal antibody, polyclonal serum, neutralization, antibody evasion

## Abstract

The SARS-CoV-2 Omicron subvariants have dominated the pandemic due to their high transmissibility and immune evasion conferred by the spike mutations. The Omicron subvariants can spread by cell-free virus infection and cell-cell fusion, the latter of which is more effective but has not been extensively investigated. In this study, we developed a simple and high-throughput assay that provides a rapid readout to quantify cell-cell fusion mediated by the SARS-CoV-2 spike proteins without using live or pseudotyped virus. This assay can be used to identify variants of concern and to screen for prophylactic and therapeutic agents. We further evaluated a panel of monoclonal antibodies (mAbs) and vaccinee sera against D614G and Omicron subvariants, finding that cell-cell fusion is substantially more resistant to mAb and serum inhibition than cell-free virus infection. Such results have important implications for the development of vaccines and antiviral antibody drugs against cell-cell fusion induced by SARS-CoV-2 spikes.

## Main text

Severe acute respiratory syndrome coronavirus 2 (SARS-CoV-2) is a novel beta-coronavirus that causes the global coronavirus disease 2019 (COVID-19) pandemic. The spike protein on the viral particles of SARS-CoV-2 mediates cell entry by binding to the human angiotensin-converting enzyme 2 (hACE2) receptor, which triggers the fusion of viral and cellular membranes (1). The spike protein is also the target for neutralizing antibodies, which are critical for the host ‘s immunity to control viral infection (2, 3).

Previous studies have shown that the SARS-CoV-2 spike protein is capable of mediating both cell-free virus infection and cell-cell transmission. Compared to cell-free virus infection, cell-cell transmission is a more efficient mode of spreading due to the direct transfer of viral particles between cells facilitated by tight cell-cell contact (4-7). Cell-cell transmission has also been observed in other coronaviruses such as the infectious bronchitis virus (IBV) and SARS-CoV (8). Like these coronaviruses, SARS-CoV-2 cell-cell transmission involves S1/S2 cleavage, cell-cell fusion, and syncytia formation (6). Notably, SARS-CoV-2 has been shown to exhibit higher cell-cell transmission activity than SARS-CoV (4).

Cell-free virus infection of Omicron subvariants, with additional spike mutations, is more resistant to antibody neutralization relative to the ancestral strain D614G (9). Whether cell-cell fusion mediated by Omicron subvariants has such antibody evasion properties, however, is not fully understood. Herein, we developed a simple, rapid, and high-throughput assay to quantitatively analyse cell-cell fusion. We further showed that cell-cell fusion induced by SARS-CoV-2 spike protein is largely resistant to the inhibition by monoclonal antibodies (mAbs) and vaccinee sera, as compared to the cell-free virus infection.

The experimental design of the cell-cell fusion assay is illustrated in **Figure 1A**. The TZM-bl-hACE2 target cell line was generated from parental TZM-bl cell line through stable transduction of hACE2 using a lentiviral system, followed by successive rounds of fluorescence-activated cell sorting (FACS) and selection of transduced cells with puromycin. Cell surface staining data revealed that TZM-bl-hACE2 cells displayed elevated levels of hACE2 compared to TZM-bl cells and Vero-E6 cells (**Figure S1A**), which are commonly used as target cells in pseudovirus assays for human immunodeficiency virus (HIV)-1 and SARS-CoV-2, respectively. The donor 293T cells were transiently transfected with HIV-1 Trans-Activator of Transcription (Tat) and spike genes, before being co-cultured with the TZM-bl-hACE2 target cells at a ratio of 2:1. Cell-cell fusion could be evaluated both qualitatively, via observation with a light microscope (**Figure S1B**), and quantitatively, through a luciferase reporter assay (**Figure S1C**). Cell-cell fusion mediated by the D614G and BA.1 variants was assessed at various time points after co-culture, and it was observed that the fusion activity of both variants increased within the first 16-24 hours of co-culture (**Figure S1C**). However, the fusion activity of D614G decreased after 24 hours, while that of BA.1 remained stable during the tested period after 16 hours of co-culture. Therefore, a co-culture time of 16 hours was chosen for subsequent experiments to ensure comparable levels of cell-cell fusion for both variants. Additionally, it was observed that D614G spike mediated slightly higher cell-cell fusion activity than BA.1 spike, despite the fact that BA.1 spike has higher affinity for ACE2 than D614G spike (10). To investigate this discrepancy, the expression and processing of the spike protein in the donor cells (293T) were examined (**Figure S2**). The results showed that the cleavage efficiency of BA.1 and other Omicron subvariants was substantially lower than that of D614G, resulting in lower levels of mature S1/S2 trimers in expressing cells, which may have contributed to the observed reduction in cell-cell fusion activity for BA.1 compared to D614G.

**Figure 1.**
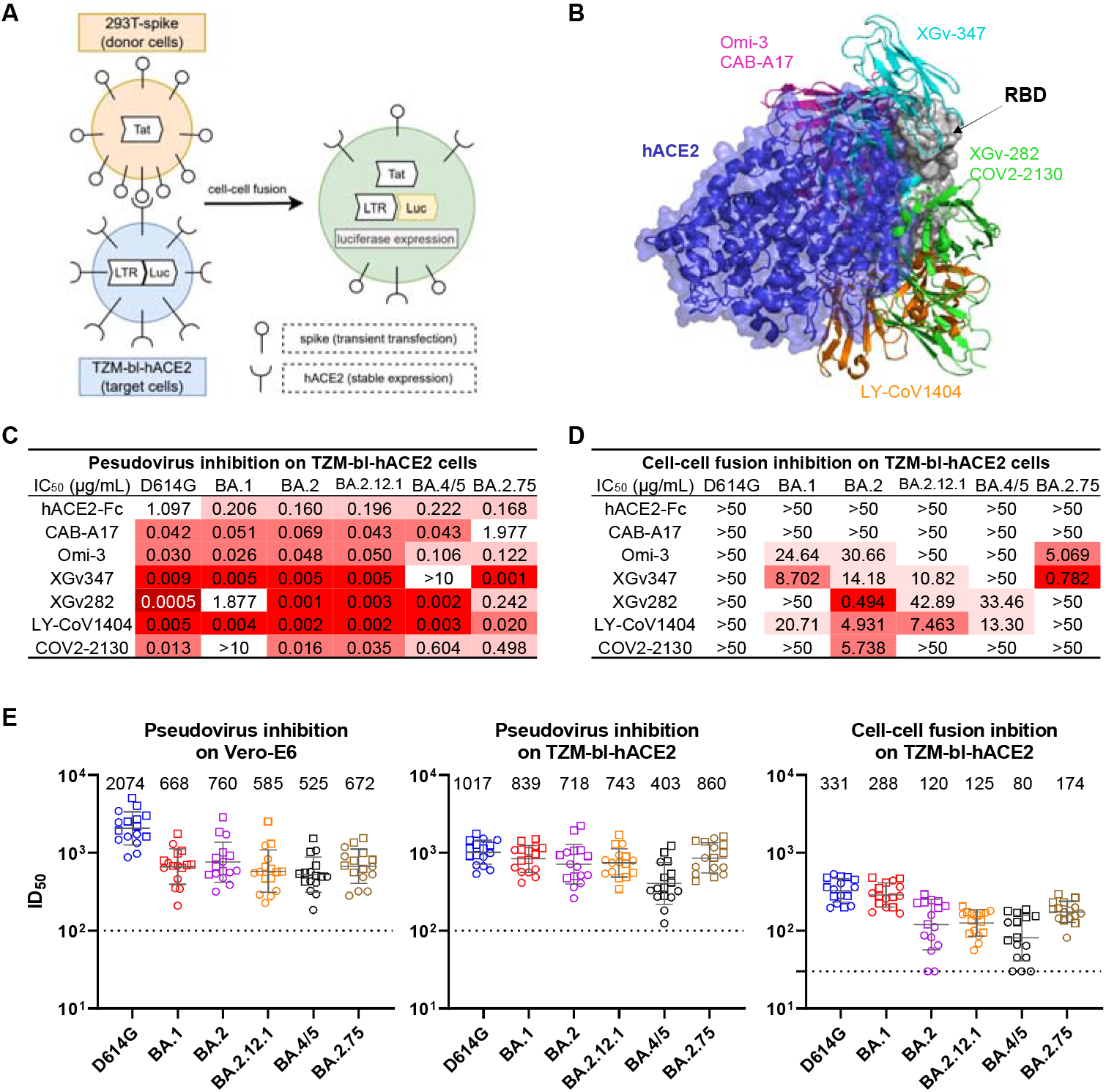
Resistance of cell-cell fusion induced by SARS-CoV-2 to monoclonal antibody and polyclonal serum neutralisation. (**A**) Schematic illustration of the cell-cell fusion assay. Briefly, 293T cells that were transiently transfected with the HIV-1 trans-activator of transcription (Tat) gene and SARS-CoV-2 spike gene were used as the donor cells. The target cells were TZM-bl-hACE2 cells that contained the firefly luciferase (luc) gene regulated by the long terminal repeat (LTR) promoter. Upon co-culture, spike protein on donor cells bound to the hACE2 receptors on target cells and initiated membrane fusion. In the syncytia, Tat proteins activated the LTR promoter, causing luc gene expression, which was quantified by luciferase reporter assay to evaluate the cell-cell fusion activity. (**B**) Structural modelling of RBD, hACE2, and mAbs. Different colors indicate different classes of mAbs, as grouped by their epitopes. (**C**) Pseudovirus neutralization IC_50_ titres for mAbs against D614G and Omicron subvariants on TZM-bl cells. (**D**) Cell-cell fusion inhibition IC_50_titres for mAbs against D614G and Omicron subvariants. (**E**) Pseudovirus neutralization and cell-cell fusion inhibition ID_50_ titres for sera against D614G and Omicron subvariants. Each point represents a serum sample, with square and circle showing samples from individuals who received 3 doses of COVID-19 mRNA vaccines with and without breakthrough infection, respectively. Data are shown as geometric mean ± SD. Horizontal solid bar indicates the geometric mean of ID_50_ titres. Horizontal dotted line denotes the limit of detection (LOD) at 100 or 30. Data shown in **C**-**E** are representative of one in three independent experiments.

We initially assessed the susceptibility of TZM-bl-hACE2 cells to SARS-CoV-2 infection using vesicular stomatitis virus (VSV)-pseudotypes. The results showed a dose-dependent infection of TZM-bl-hACE2 cells by the wild-type SARS-CoV-2 D614G, while TZM-bl cells were not infected, suggesting that hACE2 receptors were necessary for virus entry (**Figure S3A**). Furthermore, we tested the infectivity of multiple Omicron subvariants and found that they were all capable of infecting TZM-bl-hACE2 cells, although with varying levels of infectivity (**Figure S3B**). Through these evaluations, a rapid and reliable platform was developed and validated, allowing for the comparison of SARS-CoV-2 variant spikes in mediating cell-free virus infection and cell-cell fusion, as well as for testing the inhibitory effects of mAbs and vaccinee sera on these processes.

We then assessed the efficacy of mAbs and human ACE2 (hACE2) in blocking cell-free pseudovirus infection and cell-cell fusion mediated by the spikes of SARS-CoV-2 variants. The mAbs used in this study included CAB-A17 (11), Omi-3 (12), XGv347 (13), XGv282 (13), LY-CoV1404 (14), and COV2-2130 (15), which could neutralize SARS-CoV-2 by recognizing the receptor binding domain (RBD) on the spike and preventing hACE2 binding. **Figure 1B and Figure S4** display the structural modeling of the binding of these RBD-directed mAbs and hACE2 to the RBD of D614G spike. The neutralization activity of aforementioned mAbs was evaluated with VSV-pseudotyped SARS-CoV-2 on Vero-E6 cells (**Figure S5**) and TZM-bl-hACE2 cells (**Figures 1C and S6A**). Our results indicated that all mAbs effectively inhibited cell-free infection of D614G, with the 50% inhibitory concentrations (IC_50_) less than 0.3 μg/mL on TZM-bl-hACE2 cells and less than 0.1 μg/mL on Vero-E6 cells. Although the neutralizing efficacy of some mAbs was significantly impaired by Omicron, such as XGv347 against BA.4/5 and COV2-2130 against BA.1, most mAbs remained potent against the cell-free virus infection of multiple Omicron subvariants. The IC_50_ values on TZM-bl-hACE2 cells were higher than those on Vero-E6 cells, likely because TZM-bl-hACE2 cells had a higher level of hACE2 expression, which hinders the inhibition of virus entry (**Figure S1A**). As consistent with the pseudovirus neutralization data, we found that mAbs CAB-A17, Omi-3, XGv347, XGv282, and LY-CoV1404 also effectively neutralized authentic BA.1 on Vero-E6 cells with IC_50_ ranging from 0.007 to 0.217 μg/mL (**Figure S7**). But in contrast, our results showed that none of the mAbs tested could potently and broadly inhibit cell-cell fusion mediated by SARS-CoV-2 spikes (**Figures 1D and S6B**). They retained weak potency against some of the Omicron subvariants we tested. Among the variants tested, cell-cell fusion induced by D614G was the most resistant to mAb inhibition, followed by BA.4/5, while BA.2 was the most sensitive. Overall, our results demonstrated that cell-to-cell fusion mediated by SARS-CoV-2 is more resistant to mAb inhibition than cell-free virus infection.

We further profiled the inhibition efficiency of serum samples in cell-free virus infection and cell-cell fusion mediated by SARS-CoV-2 spikes. Sera were collected from a study cohort in which each subject had received three doses of mRNA-based SARS-CoV-2 vaccines. The clinical information of this study cohort is provided in **Table S1**. In both Vero-E6 cells and TZM-bl-hACE2 cells, VSV-pseudotyped SARS-CoV-2 variants were sensitive to serum neutralization, with the 50% inhibitory dose (ID_50_) titers ranging from 525 to 2074 on Vero-E6 cells, and from 403 to 1017 on TZM-bl-hACE2 cells (**Figures 1E and S8**). The ID_50_ titers on the TZM-bl-hACE2 cells were slightly lower than those on the Vero-E6 cells, which is consistent with what was observed in the pseudovirus neutralization assay to mAbs. However, in the cell-cell fusion inhibition assay on TZM-bl-hACE2 cells, the ID50 titers to serum inhibition were more than 3-fold lower than those obtained from the cell-free virus neutralization assay on the same cell line. Our data suggest that cell-cell fusion induced by SARS-CoV-2 spike protein is more resistant to inhibition by vaccinee sera, compared to cell-free virus infection mediated by the spike protein on viral particles. Among the Omicron subvariants we tested, the cell-cell fusion mediated by BA.4/5 showed the strongest evasion to vaccinee sera.

It was previously shown that cell-cell transmission of SARS-CoV-2 pre-Omicron variants is more efficient and more refractory to blockade by mAbs and convalescent sera than cell-free infection (4). In this study, we further demonstrate that the cell-cell fusion mediated by various Omicron subvariants is also resistant to inhibition by mAbs (**Figure 1D**) and vaccinee sera (**Figure 1E**). Future prophylactic or therapeutic agents should consider strategies to suppress the cell-cell fusion of SARS-CoV-2.

It should be noted that all experiments in this study were performed *in vitro*, and it is possible that the results may differ from *in vivo* outcomes. In addition, only humoral immunity against cell-cell transmission of SARS-CoV-2 was evaluated, and the role of cellular immunity requires further investigation. Lastly, cell-cell transmission was not measured directly, but indirectly through a cell-cell fusion assay. If SARS-CoV-2 triggers intense cell-cell fusion, giant syncytia may form and result in cell death, which in turn lessens cell-cell transmission.

In summary, we designed and validated a rapid and reliable cell-cell fusion assay for SARS-CoV-2. Using this assay, we demonstrated that mAbs and vaccinee sera could not effectively inhibit the cell-cell fusion mediated by SARS-CoV-2 spikes, in contrast to cell-free infection. Such results suggest that developing vaccines and antiviral antibody drugs that target the cell-cell fusion of SARS-CoV-2 may be a promising approach.

## Acknowledgements

We thank C. Lu and the Columbia Center for Translational Immunology (CCTI) Flow Cytometry Core for their help with cell sorting.

