## Supplemental information for "Impaired potency of neutralizing antibodies against cell-cell fusion mediated by SARS-CoV-2"

### Materials and methods

#### *Vaccinee sera*

Serum specimens were obtained from subjects who received a three-dose regimen of either mRNA-1273 or BNT162b2 vaccine, through a collection process performed at Columbia University Irving Medical Center. The collection was conducted after obtaining written informed consent from each participant, in accordance with protocols approved by the Institutional Review Board of Columbia University. The relevant clinical information is provided in **Table S1**.

#### *Cell lines*

Expi293 cells (A14527) were procured from Thermo Fisher Scientific and maintained in Expi293$^{TM}$ Expression Medium with 0.5% penicillin-streptomycin (Thermo Fisher Scientific). These cells were grown at 37°C in an 8% $CO_2$ atmosphere with 125 rpm shaking. 293T cells (CRL-3216) and Vero-E6 cells (CRL-1586) were obtained from the American Type Culture Collection (ATCC), and TZM-bl cells (ARP-8129) were obtained from the NIH HIV Reagent Program. These cells were maintained in Dulbecco modified Eagle medium (DMEM) with 10% fetal bovine serum (FBS) and 1% penicillin-streptomycin, and were grown at 37°C under 5% $CO_2$ conditions.

The TZM-bl-hACE2 cells were generated through lentiviral introduction of the human ACE2 gene into TZM-bl cells. The procedure involved cloning the full-length human ACE2 gene into the pLVX-IRES-Puro plasmid, co-transfecting 293T cells with the pMD2.G and psPAX2 plasmids to produce the lentivirus containing the hACE2 gene, infecting TZM-bl cells with the lentivirus, and culturing the infected cells in the presence of puromycin. The cells were then isolated by fluorescence-activated cell sorting (FACS, BD FACSAria III; BD Biosciences) to select for cells expressing high and uniform levels of hACE2. The TZM-bl-hACE2 cells were maintained in DMEM supplemented with 10% FBS, 1% penicillin-streptomycin, and 10 μg/ml of puromycin, and were incubated at 37°C under 5% CO2.

All cell lines tested mycoplasma negative and had normal morphology under light microscope before experimental use.

#### *Monoclonal antibodies (mAbs) and hACE2*

In-house expression of antibodies was accomplished by transfecting Expi293 cells (Thermo Fisher Scientific) with both the heavy and light chains of each antibody, using 1 mg/mL polyethylenimine (PEI) as the transfection reagent. On the fourth day after transfection, affinity purification was performed from the cellular supernatants using rProtein A Sepharose (GE). To obtain hACE2-Fc, the ectodomain of human ACE2, fused with a Fc tag at its C-terminus, was expressed by transfecting the plasmid pcDNA3-sACE2-WT (732)-IgG1 (Addgene plasmid #154104) into Expi293 cells using PEI. The secreted protein was purified using rProtein A Sepharose.

The size and purity of all proteins were verified using sodium dodecyl-sulfate polyacrylamide gel electrophoresis (SDS-PAGE) prior to experimentation.

#### *Pseudovirus production*

Pseudotyped SARS-CoV-2 were generated by incorporating the spike glycoprotein of SARS-CoV-2 into vesicular stomatitis virus (VSV). This process involved transfecting 293T cells with a construct encoding the SARS-CoV-2 spike glycoprotein using PEI, and then infecting the transfected cells with the ΔG-luciferase-expressing VSV (G*ΔG-luciferase) from Kerafast Inc. The infection was carried out 24 hours post-transfection and after 2 hours of infection, the cells were washed and changed to fresh medium. Subsequently, the culture was continued for an additional 24 hours, after which the supernatant was harvested, clarified via centrifugation, and stored at -80°C.

***Western blot***

293T cells were transfected to produce VSV-based particles pseudotyped with SARS-CoV-2 spikes, as described above. In parallel with harvesting the pseudoviruses from the cell supernatants, 293T cells were washed with PBS for 3 times and lysed using 1% NP-40 at 4˚C for 10 min. Cell lysates were then clarified by high-speed centrifugation (18,000g for 10 min) and analyzed by Western blotting with the following primary antibodies: rabbit anti-SARS-Spike S1 (Sino Biological), rabbit anti-SARS-Spike S2 (Sino Biological), and mouse anti-VSV NP (Millipore). The Western blots were developed with the following secondary antibodies: HRP-conjugated anti-rabbit antibody (Cytiva) or HRP-conjugated goat anti-mouse antibody (Jackson ImmunoResearch).

***Pseudovirus neutralization assay***

The viral titers of all pseudoviruses were first adjusted to ensure consistent viral input in each assay. Heat-inactivated sera, monoclonal antibodies, and hACE2 were prepared in media and dispensed into 96-well plates in triplicate, with serial dilutions starting at 1:100 for sera and 10 μg/mL for antibodies and hACE2-Fc. The prepared pseudoviruses were then mixed with the samples, incubated at 37°C for 1 hour, and virus-only wells were included as a control. Note that no incubation was performed for hACE2. $3 \times 10^4$ cells per well were added and the mixture was incubated for 16 hours at 37°C. Subsequently, cells were lysed and luciferase activity was measured using the Luciferase Assay System (Promega) and SoftMax Pro v.7.0.2 (Molecular Devices) as per the manufacturer's instructions. Neutralization curves, $IC_{50}$ titers, and $ID_{50}$ titers were determined by fitting a non-linear five-parameter dose-response curve to the data using Prism v.9.2 (GraphPad).

***Authentic virus neutralization assay***

The BA.1 authentic virus was obtained from BEI (NR-56481, hCoV-19/USA/GA-EHC-2811C/2021). Vero-E6 cells were plated at $1.5 \times 10^4$ cells/well in DMEM supplemented with 10% FCS and 1× Penicillin/Streptomycin (ThermoFisher). Next day, authentic BA.1 virus (100 $TCID_{50}$) was mixed with the 5-fold dilution series of antibodies for 1 hour at 37 °C. Then the virus-antibody mixture was inoculated onto the 96 well plates in quadruplicate and cultured at 37 °C with 5% $CO_2$. On day 3, the CPE was scored (from 0 to 4+) under the inverted microscopy and $IC_{50}$ values were estimated using GraphPad Prism v.9.2.

***Flow cytometry***

The cells were trypsinized and suspended in a FACS buffer consisting of PBS and 2% FBS. Subsequently, the cells were incubated with anti-human ACE2 antibody (Biolegend, Cat# 503602) at 4°C for 1 hour, followed by the incubation of APC anti-human IgG Fc recombinant second antibody (Biolegend, Cat# 366906) at 4°C for an additional hour. The cells were then resuspended in FACS buffer, and the expression level of ACE2 was quantified using an LSRII flow cytometer (BD Biosciences).

***Cell-cell fusion and inhibition assay***

Donor (293T) cells were transfected with the transcriptional activator Tat (tat) gene and the relevant spike gene and incubated at 37°C under 5% $CO_2$ for 24 hours. Then, donor cells (transfected 293T cells) and target cells (TZM-bl cells or TZM-bl-hACE2 cells) were trypsinized and co-cultured at 37°C for 16 hours in triplicate in 96-well plates at a density of $4 \times 10^4$ cells/well and $2 \times 10^4$ cells/well, respectively. Following incubation, cells were lysed, and luciferase activity was measured using the Luciferase Assay System (Promega) and SoftMax Pro v.7.0.2 (Molecular Devices) as per the manufacturer's instructions.

To measure the inhibition in cell-cell fusion, heat-inactivated sera, mAbs, or hACE2 were prepared in culture medium and added in triplicate to 96-well plates, with serial dilutions starting at 1:30 for sera and 50 μg/mL for antibodies and hACE2-Fc. Donor cells were then added to the wells at a density of $4 \times 10^4$ cells per well and incubated at 37°C for 1 hour. Then, TZM-bl-hACE2 cells were added to the wells at a density of $2 \times 10^4$ cells per well and incubated at 37°C for another 16 hours. Control wells containing only donor cells were included on all plates. Neutralization curves, $IC_{50}$ titers, and $ID_{50}$ titers were generated by fitting a non-linear five-parameter dose-response curve to the data using Prism v.9.2 (GraphPad).

***Structural modeling of RBD mutations***

The structures of antibody-spike complexes were obtained from Protein Data Bank (PDB): ACE2, 7C8D; Omi-3, 7ZF3; XGv347, 7WED; XGv282, 7WLC; LY-CoV1404, 7MMO. Structural modeling was generated using PyMOL v.2.3.2 (Schrodinger, LLC).

**Table S1. Demographics of clinical cohorts.**

| Sample ID | Vaccine type and infected strain | Days post-vaccination or *infection (after last exposure) | Documented COVID-19 | Age | Gender |
|---|---|---|---|---|---|
| Vaccination samples | | | | | |
| Q1 | mRNA-1273/mRNA-1273/mRNA-1273 | 29 | No | 66 | Female |
| Q2 | BNT162b2/BNT162b2/BNT162b2 | 30 | No | 68 | Male |
| Q3 | BNT162b2/BNT162b2/BNT162b2 | 14 | No | 64 | Female |
| Q4 | BNT162b2/BNT162b2/BNT162b2 | 34 | No | 55 | Male |
| Q5 | BNT162b2/BNT162b2/BNT162b2 | 34 | No | 45 | Male |
| Q6 | BNT162b2/BNT162b2/BNT162b2 | 15 | No | 50 | Female |
| Q7 | BNT162b2/BNT162b2/BNT162b2 | 15 | No | 48 | Female |
| Q8 | BNT162b2/BNT162b2/BNT162b2 | 29 | No | 71 | Male |
| Breakthrough samples | | | | | |
| Q36 | BNT162b2/BNT162b2/BNT162b2/Ad26.COV2.S/BA.2 | *22 | Yes | 69 | Male |
| Q50 | mRNA-1273/mRNA-1273/mRNA-1273/BA.2 | *14 | Yes | 34 | Male |
| Q51 | BNT162b2/BNT162b2/mRNA-1273/BA.2 | *19 | Yes | 33 | Female |
| Q52 | BNT162b2/BNT162b2/mRNA-1273/BA.2 | *18 | Yes | 29 | Female |
| Q55 | BNT162b2/BNT162b2/mRNA-1273/BA.2 | *18 | Yes | 41 | Female |
| Q56 | mRNA-1273/mRNA-1273/mRNA-1273/BA.2 | *21 | Yes | 36 | Female |
| Q57 | BNT162b2/BNT162b2/mRNA-1273/BA.2 | *32 | Yes | 28 | Male |
| Q58 | BNT162b2/BNT162b2/mRNA-1273/BA.2 | *23 | Yes | 33 | Female |
| Healthy samples | | | | | |
| ADARC13 | Unvaccinated healthy donor | N/A | No | 32 | Female |
| ADARC14 | Unvaccinated healthy donor | N/A | No | 46 | Male |

N/A, not available.

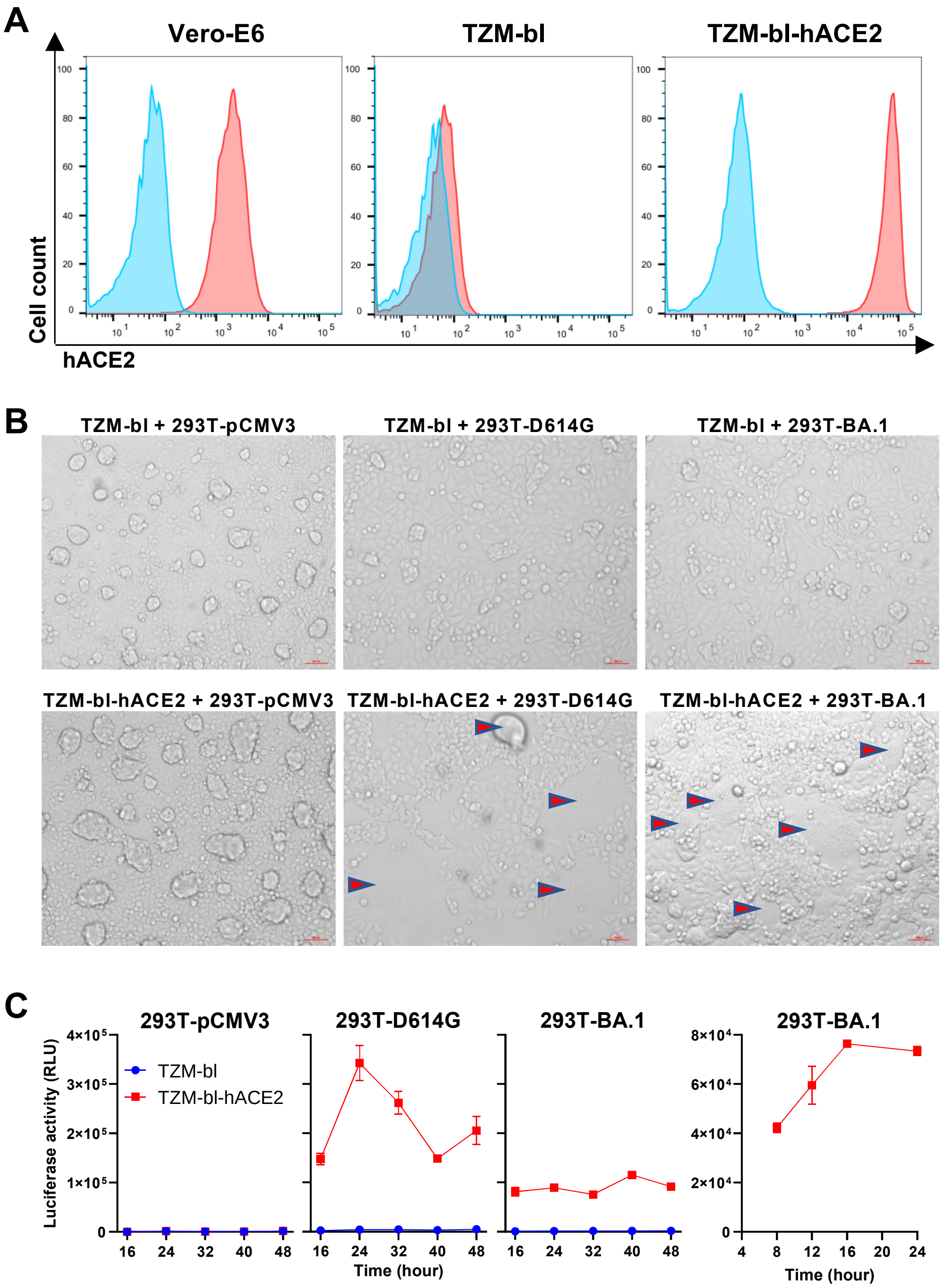


**Figure S1. Development and validation of the cell-cell fusion assay.**

A. The expression levels of hACE2 on the surfaces of Vero-E6, TZM-bl, and TZM-bl-hACE2 cells were determined by FACS analysis. The color blue indicates unstained controls; red indicates the hACE2 expression levels.

B. Microscopic examination was conducted to observe the cell-cell fusion under various conditions. The term "293T-pCMV3" refers to 293T cells transfected with the pCMV3 vector but without the SARS-CoV-2 spike. The fused cells are indicated by arrowheads.

C. The efficiency of cell-cell fusion mediated by the D614G and BA.1 spikes was determined at various time points after co-culture, as measured by luciferase reporter assay in the syncytia. Data shown in **A-C** are representative of one in three independent experiments.

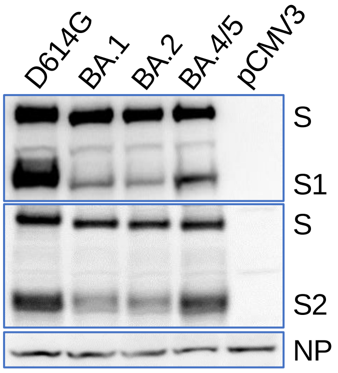


**Figure S2. Expression and processing of SARS-CoV-2 D614G and Omicron subvariants.** 293T cells producing VSV particles pseudotyped with the indicated SARS-CoV-2 spike protein were used to prepare cell lysates, which were western blotted and probed with rabbit anti-S1, anti-S2, and anti-VSV NP antibodies. Cells transfected with the pCMV3 vector were used as a negative control. Data shown are representative of one in three independent experiments.

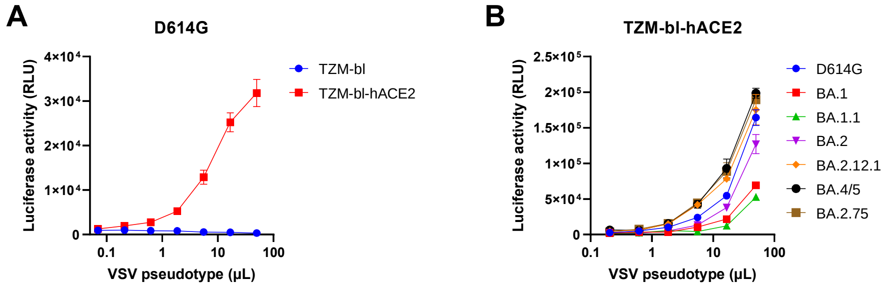


**Figure S3. Susceptibility of TZM-bl-hACE2 cells to infection by VSV-pseudotyped SARS-CoV-2 variants.**

A. Infectivity of pseudotyped D614G in TZM-bl and TZM-bl-hACE2 cells.

B. Infectivity of pseudotyped Omicron subvariants in TZM-bl-hACE2 cells.

Data shown in **A-B** are representative of one in three independent experiments.

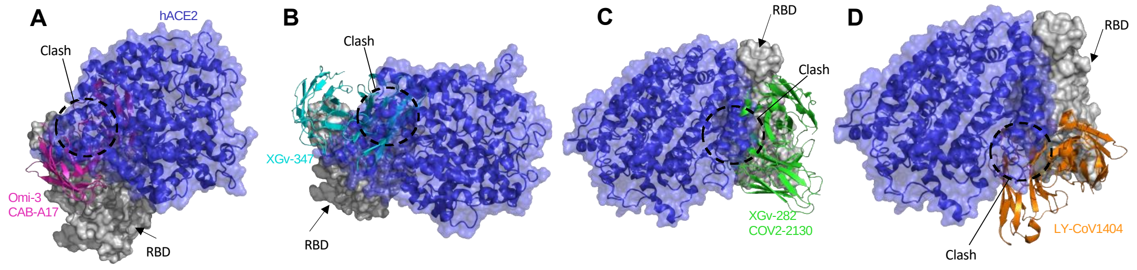


**Figure S4. Structural modeling of RBD-directed mAbs Omi-3 and CAB-A17 (A), XGv-347 (B), XGV-282 and COV2-2130 (C), and LY-CoV1404 (D) in complex with RBD and hACE2.**

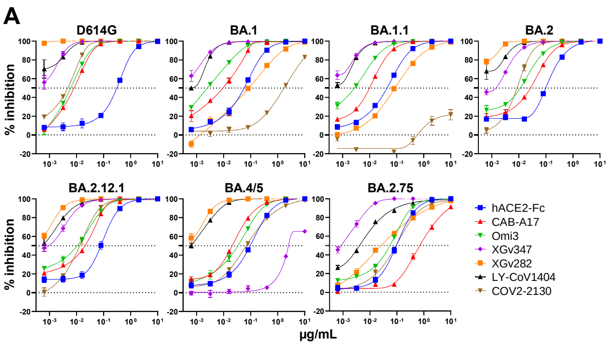


**B**

| Vero-E6 | | | | | | | |
|---|---|---|---|---|---|---|---|
| $IC_{50}$ (μg/mL) | D614G | BA.1 | BA.1.1 | BA.2 | BA.2.12.1 | BA.4/5 | BA.2.75 |
| hACE2-Fc | 0.309 | 0.057 | 0.044 | 0.098 | 0.079 | 0.096 | 0.096 |
| CAB-A17 | 0.008 | 0.009 | 0.010 | 0.027 | 0.021 | 0.021 | 0.714 |
| Omi3 | 0.006 | 0.003 | 0.003 | 0.011 | 0.011 | 0.034 | 0.055 |
| XGv347 | 0.0001 | <0.001 | <0.001 | 0.001 | 0.001 | 2.505 | 0.001 |
| XGv282 | <0.001 | 0.084 | 0.086 | <0.001 | <0.001 | <0.001 | 0.026 |
| LY-CoV1404 | <0.001 | 0.001 | <0.001 | <0.001 | 0.001 | 0.001 | 0.004 |
| COV2-2130 | 0.005 | 1.721 | >10 | 0.009 | 0.013 | 0.069 | 0.094 |

**Figure S5. Neutralization sensitivity of monoclonal antibodies (mAbs) against VSV-pseudotyped SARS-CoV-2 variants on Vero-E6 cells.**

A. Neutralization curves. Horizontal dotted line indicates 50% inhibition.

B. Neutralization $IC_{50}$ titers. The darker colors indicate antibodies with higher neutralization activities. Data shown in **A-B** are representative of one in three independent experiments.

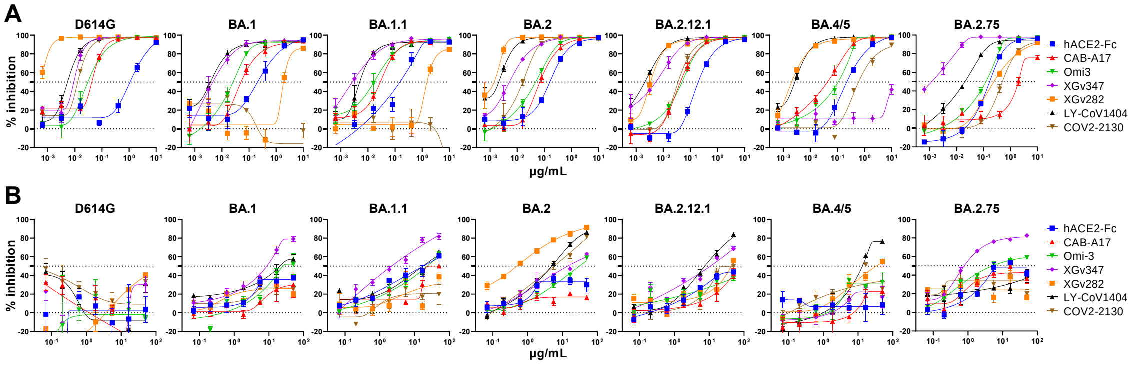


**Figure S6. Neutralization curves of mAbs against the VSV-pseudotyped SARS-CoV-2 variants (A) and inhibition ability of mAbs in the cell-cell fusion mediated by the SARS-CoV-2 spikes (B) on TZM-bl-hACE2 cells.** Horizontal dotted line indicates 50% neutralization or inhibition. Data shown in **A-B** are representative of one in three independent experiments.

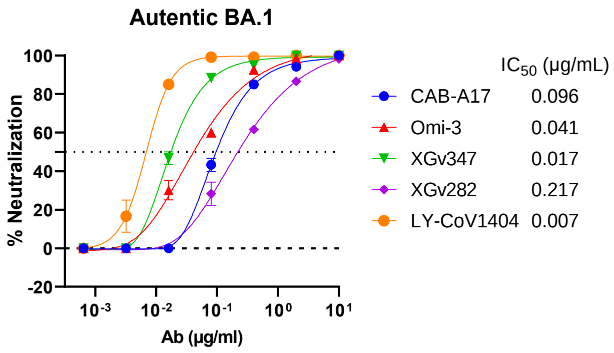


**Figure S7. Neutralization activity of mAbs against authentic BA.1 on Vero-E6 cells.** Data are shown as mean ± SEM and $IC_{50}$ values for mAbs are presented.

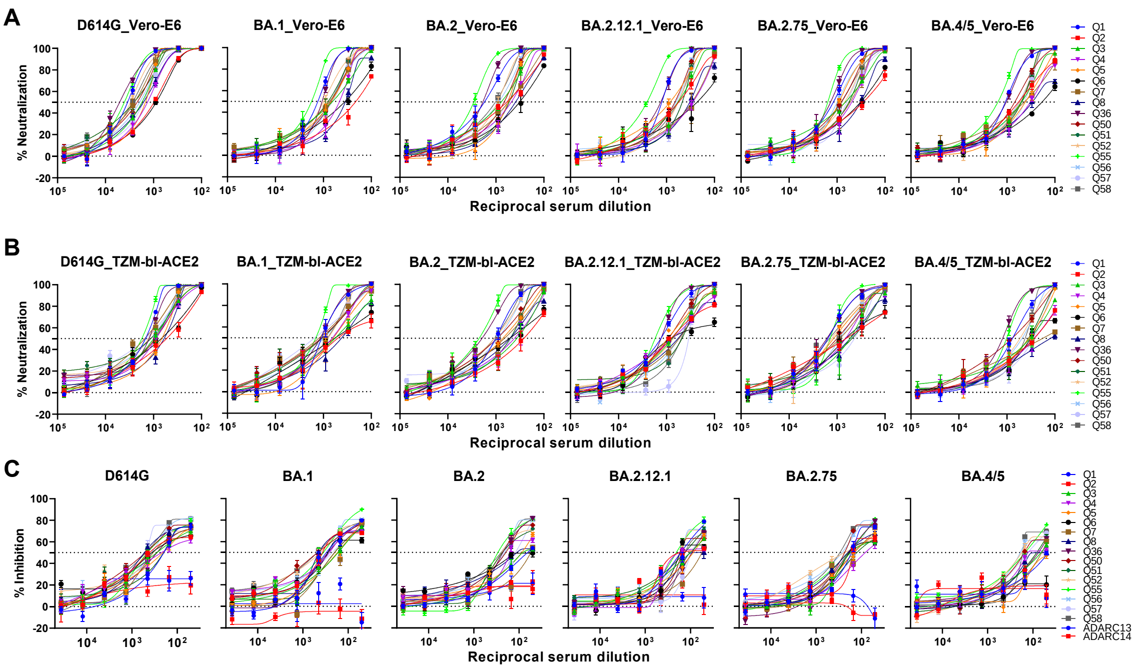


**Figure S8. Neutralization activity of the serum samples from healthy donors who received three doses of mRNA vaccines against (A) pseudovirus infection in Vero-E6 cells, (B) pseudovirus infection in TZM-bl-hACE2 cells, and (C) cell-cell fusion in the TZM-bl-hACE2 target cells**. Horizontal dotted lines indicate 0% and 50% inhibition levels. Data shown in **A-C** are representative of one in three independent experiments.
